# Protective face mask filter capable of inactivating SARS-CoV-2, and methicillin-resistant *Staphylococcus aureus* and *Staphylococcus epidermidis*

**DOI:** 10.1101/2020.11.24.396028

**Authors:** Miguel Martí, Alberto Tuñón-Molina, Finn Lillelund Aachmann, Yukiko Muramoto, Takeshi Noda, Kazuo Takayama, Ángel Serrano-Aroca

**Affiliations:** Biomaterials and Bioengineering Lab, Centro de Investigación Traslacional San Alberto Magno, Universidad Católica de Valencia San Vicente Mártir, c/Guillem de Castro 94, Valencia 46001, Spain; NOBIPOL, Department of Biotechnology and Food Science, NTNU Norwegian University of Science and Technology, Sem Sælands vei 6-8, N-7491 Trondheim, Norway; Laboratory of Ultrastructural Virology, Institute for Frontier Life and Medical Sciences, Kyoto University, Kyoto 606-8507, Japan; Center for iPS Cell Research and Application, Kyoto University, Kyoto 606-8397, Japan

**Keywords:** SARS-CoV-2, MRSA, MRSE, face mask filter, benzalkonium chloride, COVID-19, multidrug-resistant bacteria

## Abstract

Face masks have globally been accepted to be an effective protective tool to prevent bacterial and viral transmission, especially against indoor aerosol transmission. However, commercial face masks contain filters that are made of materials that are not capable of inactivating neither SARS-CoV-2 nor multidrug-resistant bacteria. Therefore, symptomatic and asymptomatic individuals can infect other people even if they wear them because some viable viral or bacterial loads can escape from the masks. Furthermore, viral or bacterial contact transmission can occur after touching the mask, which constitutes an increasing source of contaminated biological waste. Additionally, bacterial pathogens contribute to the SARS-CoV-2 mediated pneumonia disease complex and their resistance to antibiotics in pneumonia treatment is increasing at an alarming rate. In this regard, herein, we report the development of a novel protective non-woven face mask filter fabricated with a biofunctional coating of benzalkonium chloride that is capable of inactivating SARS-CoV-2 in one minute of contact, and the life-threatening methicillin-resistant *Staphylococcus aureus* and *Staphylococcus epidermidis.* Nonetheless, despite the results obtained, further studies are needed to ensure the safety and correct use of this technology for the mass production and commercialization of this broad-spectrum antimicrobial face mask filter. Our novel protective non-woven face mask filter would be useful for many health care workers and researchers working in this urgent and challenging field.

## 1. Introduction

The Severe Acute Respiratory Syndrome Coronavirus 2 (SARS-CoV-2) was first reported in Wuhan, Hubei province, China, in December 2019^1^. The rapid spread of this pathogen, which has caused the current COVID-19 pandemic, is putting at high risk the health and economy of the most developed and underdeveloped countries. According to the World Health Organisation (WHO), the current COVID-19 outbreak has 59,342,857 global cases and 1,399,373 global deaths in more than 200 countries (data of 24^th^ November 2020)^2^. SARS-CoV-2 is the third coronavirus causing severe pneumonia^3,4^, an infection of the lungs caused usually by bacteria and viruses^5,6^. The death risk of viral pneumonias can increase when co-infection can be caused by viruses in the setting of community-acquired bacterial pneumonia such as the lethal *Streptococcus pneumoniae*^7–10^, with additional symptoms of bacterial pneumonia^11^. New pathogens, such as SARS-CoV-2, that can coexist with a broad range of other types of clinically relevant bacteria, including multidrug-resistant strains, constitutes a real life-threatening to humans during the approaching cold season. In addition, antibiotic resistance in bacterial pneumonia treatment is a wide-spread problem that is increasing at an alarming rate^12,13^. The SARS-CoV-2 pathogen is stable from hours to days in aerosols and surfaces of different chemical nature such as copper, cardboard, plastic, aluminium, or stainless steel surfaces, demonstrating that infections can be easily transmitted through the air *via* the microdroplets or direct contact after touching contaminated surfaces^14–18^. This coronavirus can spread faster than its two ancestors SARS-CoV and MERS-CoV^19^ through coughing, sneezing, touching or breathing ^20^, and more broadly through asymptomatic carriers^21–23^. Recent studies have demonstrated that direct indoor aerosol transmission or via ventilation systems can be potentially transmit SARS-CoV-2^24–26^. Although the confinements conducted in many countries flattened the epidemic curve before the hot season^27,28^, SARS-CoV-2 continues to spread globally. SARS-CoV-2 is an enveloped positive-sense single-stranded RNA virus^29^ that belongs to the IV Baltimore group^30^. Other enveloped RNA viruses such as influenza A (H1N1) can be inactivated by quaternary ammonium compounds such as benzalkonium chloride (BAK)^31^. It has been recently remarked, however, that further evaluation of the effectiveness of BAK against coronaviruses is needed^32^ because the Centers for Disease Control and Prevention have reported that available evidence indicates BAK has less reliable activity against certain bacteria and viruses than either of the alcohols^33^. However, a recent report has shown the *in vitro* virucidal activity of ethanol (70%), povidone-iodine (7.5%), chloroxylenol (0.05%), chlorhexidine (0.05%), or benzalkonium chloride (0.1%) was similar when used as disinfectants against SARS-CoV-2^18^. Thus, the oral rinse Dequonal, which contains BAK, has shown virucidal activity against SARS-CoV-2 under conditions mimicking nasopharyngeal secretions to support the idea that oral rinsing might reduce the viral load of saliva and could thus lower the transmission of SARS-CoV-2^34^. Furthermore, very recently, a preprint has reported an oil-in-water nanoemulsion formulation containing 0.13% BAK that has demonstrated safe and broad antiviral activity against enveloped viruses such as SARS-CoV-2, human coronavirus, respiratory syncytial virus and influenza B^35^. In that study, the repeated application of this BAK-containing nanoemulsion, twice daily for 2 weeks on to rabbit nostrils indicated safety with no irritation. In fact, this chemical compound is widely used as a disinfectant against bacteria, viruses, pathogenic fungi and mycobacteria and it has been approved by the Food and Drug Administration as a skin disinfectant^36^. In this study, we hypothesise here that the physical adsorption of BAK by the dip coating method^37^ onto the surface of a commercial non-woven fabric filter, which are commonly used in the production of face masks in the present pandemic, could produce antiviral filters that can inhibit the infection capacity of SARS-CoV-2. Non-woven filters are lightweight, flexible, resilient, provides good bacteria filtration and air permeability, and are costeffective materials for masks, has a lower manufacturing cost, are hygienic and clean as they are for single use^38^. Face masks have been accepted as effective protective tools by blocking the pass of viral and bacterial particles^39^. However, if the filters that contain the face masks are made by composite materials with antimicrobial activity, the protection of these tools could increase even more. Furthermore, due to the previously reported antibacterial activity of BAK against Gram-positive bacteria^40^, we also expect that the developed BAK filter will be able to inhibit the bacterial grow of two clinically relevant multidrug resistant bacteria: methicillin-resistant *Staphylococcus aureus* (MRSA) and *Staphylococcus epidermidis* (MRSE). In addition to the current COVID-19 pandemic, antibiotic resistance is another increasing challenge of the present century. According to the World Health Organization (WHO), antibiotic resistance will be one of the leading causes of death over other important diseases such as cancer by the year 2050^41^. In fact, MRSE is a nosocomial pathogen that is spreading globally and is often cause of catheter-associated disease, especially among low birth weight prematures^42,43^. MRSA is causing global health problems specially in medical instruments and catheters because *S. aureus* is a human pathogen that can easily develop resistance to antibiotics^44,45^. Finally, in this study, we attempted to develop a protective face mask filter capable of inactivating SARS-CoV-2, and MRSA and MRSE.

## 2. Materials and methods

### 2.1. Dip coating of commercial face filter masks

Disks specimens of approximately 10 mm in diameter were prepared with a non-woven spunlace fabric filter from NV EVOLUTIA (commercial filters used for face masks) by dry cutting with a cylindrical punch. Face mask filter (BAK Filter) disks *(n=6)* were produced by the dip coating method^37^ using commercial 70% ethyl alcohol with 0.1% *w/w* benzalkonium chloride from Montplet (Barcelona, Spain) for 1 minute at 25°C to achieve a dry BAK content, determined gravimetrically, of 0.46±0.13% *w/w.* Another face mask filter (S Filter) disks (*n=6*) were subjected to the same dip coating treatment but using only an absolute ethanol/distilled water solution (70/30% *v/v)* without BAK for 1 minute at 25°C. Untreated face mask filter (U Filter) disks (*n=6*) were produced as reference material. The disks were subsequently dried at 60°C for 48 hours to constant weight and sterilized by UV radiation for one hour per each side.

### 2.2. Characterization of the benzalkonium chloride

Nuclear magnetic resonance (NMR) was applied for the characterization of the benzalkonium chloride used in the biofunctional coating of the commercial nonwoven filters. Prior to NMR sample preparation, the ethanol/water solvent was evaporated from commercial Montplet 70% ethyl alcohol with benzalkonium chloride (99.9/0.1% *w/w)* at 25°C. After that, the sample of benzalkonium chloride was prepared by dissolving 10 mg in 550 μL D2O (D, 99.9 %) (Sigma-Aldrich, Norway) and transferred to 5mm LabScape Stream NMR tube. The NMR experiments were recorded on a BRUKER AVIIIHD 800 MHz (Bruker BioSpin AG, Fälladen, Switzerland) equipped with 5mm with cryogenic CP-TCI. All NMR recording was performed at 25°C or 37°C. For the characterization of benzalkonium chloride the following spectra were recorded: 1D proton, 2D double quantum filtered correlation spectroscopy (DQF-COSY) and 2D ^13^C heteronuclear single quantum coherence (HSQC) with multiplicity editing. TMS was used as chemical shift reference for proton and carbon chemical shifts. The spectra were recorded, processed and analysed using TopSpin 3.7 software (Bruker BioSpin).

### 2.3. Electron microscopy

A Zeiss Ultra 55 field emission scanning electron microscope (FESEM, Zeiss Ultra 55 Model) was operated at an accelerating voltage of 3 kV to observe the porous morphology of the treated and untreated non-woven face mask filters at a magnification of x100 and x1000. The filter samples were prepared to be conductive by platinum coating with a sputter coating unit.

### 2.4. Phage phi6 host culture

*Pseudomonas syringae* (DSM 21482) from the Leibniz Institute DSMZ-German collection of microorganisms and cell cultures GmbH (Braunschweig, Germany), was cultured in solid Tryptic Soy Agar (TSA, Liofilchem) and subsequently in Liquid Tryptic Soy Broth (TSB, Liofilchem). Liquid incubation was carried out at 25°C and 120rpm.

### 2.5. Phage phi 6 propagation

Phage phi 6 (DSM 21518) propagation was carried out according to the specifications provided by the Leibniz Institute DSMZ-German collection of microorganisms and cell cultures GmbH (Braunschweig, Germany).

### 2.6. Antiviral test using the biosafe viral model

A volume of 50 μL of a phage suspension in TSB was added to each filter at a titer of about 1×10^6^ plaque-forming units per mL (PFU/mL) and allowed to incubate for 1, 10 and 30 minutes. Each filter was placed in a falcon tube with 10mL TSB and sonicated for 5 minutes at 24 ° C. After that, each tube was vortexed for 1 minute. Serial dilutions of each falcon sample were made for phage titration and 100 μL of each phage dilution were contacted with 100 μL of the host strain at OD600 nm = 0.5. The infective capacity of the phage was measured based on the double-layer method^46^ where 4 mL of top agar (TSB + 0.75% bacteriological agar, Scharlau) 5 mM CaCl2 was added to the phage-bacteria mixture which was poured on TSA plates. The plates were incubated for 24-48 hours in an oven at 25°C. The phage titer of each type of sample was calculated in PFU/mL and compared with the control, that is, 50 μL of phage added to the bacteria without being in contact with any filter and without being sonicated. The antiviral activity in log reductions of titers was estimated at 1, 10 and 30 minutes of contact with the virus model. It was checked that the residual amounts of disinfectants in the titrated samples did not interfere with the titration process and the sonication-vortex treatment did not affect to the infectious capacity of the phage. The antiviral tests were performed three times during two different days (*n=6*) to ensure reproducibility.

### 2.7. Antiviral tests using SARS-CoV-2

The SARS-CoV-2 strain used in this study (SARS-CoV-2/Hu/DP/Kng/19-027) was kindly gifted from Dr. Tomohiko Takasaki and Dr. Jun-Ichi Sakuragi at the Kanagawa Prefectural Institute of Public Health. The virus was plaque-purified and propagated in Vero cells. SARS-CoV-2 was stored at −80°C.

A volume of 50 μL of a virus suspension in PBS was added to each filter at a titer dose of 1.3×10^5^ TCID50/filter, and then incubated for 1 minute at room temperature. 1 ml PBS was added to each filter, and then vortexed for 5 minutes. After that, each tube was vortexed for 5 minutes at room temperature.

Viral titers were determined through median tissue culture infectious dose (TCID50) assays inside a Biosafety Level 3 laboratory at Kyoto University. Briefly, TMPRSS2/Vero cells^47^ (JCRB1818, JCRB Cell Bank), cultured with the Minimum Essential Media (MEM, Sigma-Aldrich) supplemented with 5% fetal bovine serum (FBS), 1% penicillin/streptomycin, were seeded into 96-well plates (Thermo Fisher Scientific). Samples were serially diluted 10-fold from 10^-1^ to 10^-8^ in the culture medium. Dilutions were placed onto the TMPRSS2/Vero cells in triplicate and incubated at 37°C for 96 hours. Cytopathic effect was evaluated under a microscope. TCID50/mL was calculated using the Reed-Muench method.

### 2.8. Antibacterial tests

The agar disk diffusion tests were performed to analyse the antibacterial activity of the calcium alginate/CNFs composite films^48,49^. Lawns of methicillin-resistant *Staphylococcus aureus,* COL ^50^, and the methicillin-resistant *Staphylococcus epidermidis,* RP62A ^51^, in a concentration about 1.5 × 10^8^ CFU/mL in tryptic soy broth were cultivated on trypticase soy agar plates. The sterilized disks were placed upon the lawns of bacteria to be incubated aerobically at 37 °C for 24 hours. The antibacterial activity of the tested filter disks was expressed according to equation (1)^48^:

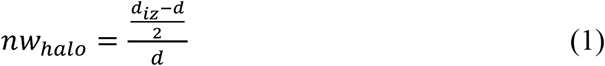

where *nw_haio_* indicates the normalised width of the antimicrobial inhibition zone, *diz* is the inhibition zone diameter and *d* referrers to the sample disk diameter. These diameters were measured by image software analysis (Image J). The tests were performed six times in different days to ensure reproducibility.

### 2.9. Statistical analysis

The statistical analyses were performed by ANOVA followed by Tukey’s posthoc test (*p > 0.05, ***p > 0.001) on GraphPad Prism 6 software.

## 3. Results and discussion

### 3.1. Nuclear magnetic resonance of benzalkonium chloride

The benzalkonium chloride used in this study for the treatment of the unwoven filters analysed by NMR is shown in Figure 1.

**Figure 1.**
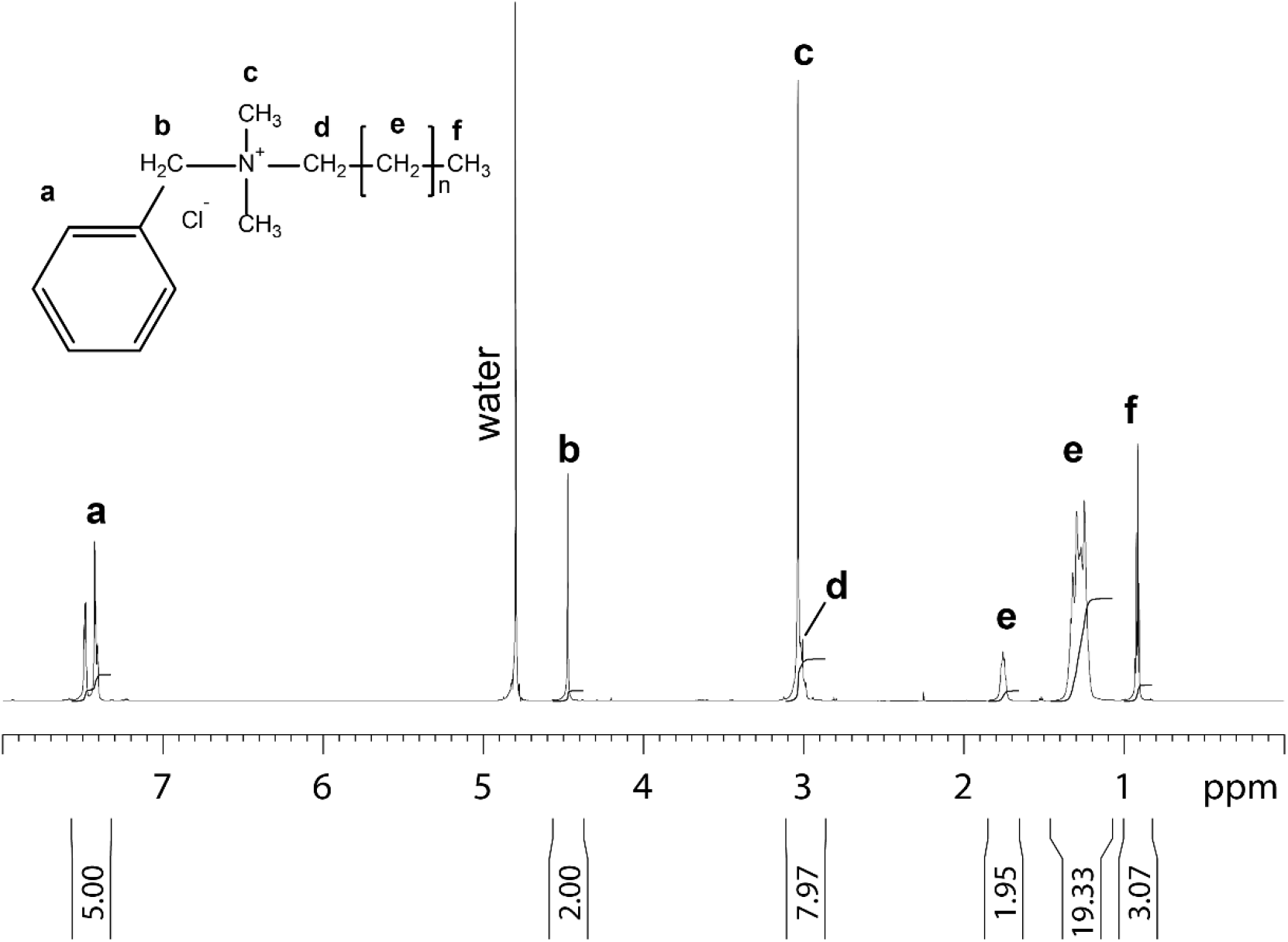
1D proton NMR spectrum of benzalkonium chloride dissolved in 99.9% D2O recorded at 25°C. Molecular structure, assignment and integral for benzalkonium chloride is shown. The letters at the molecular structure and the spectrum indicates the proton in the different chemical subgroups of benzalkonium chloride.

### 3.2. Porous morphology of the non-woven face mask filters

The porous morphology images of the commercial and treated non-woven face mask filter are shown in Figure 2.

**Figure 2.**
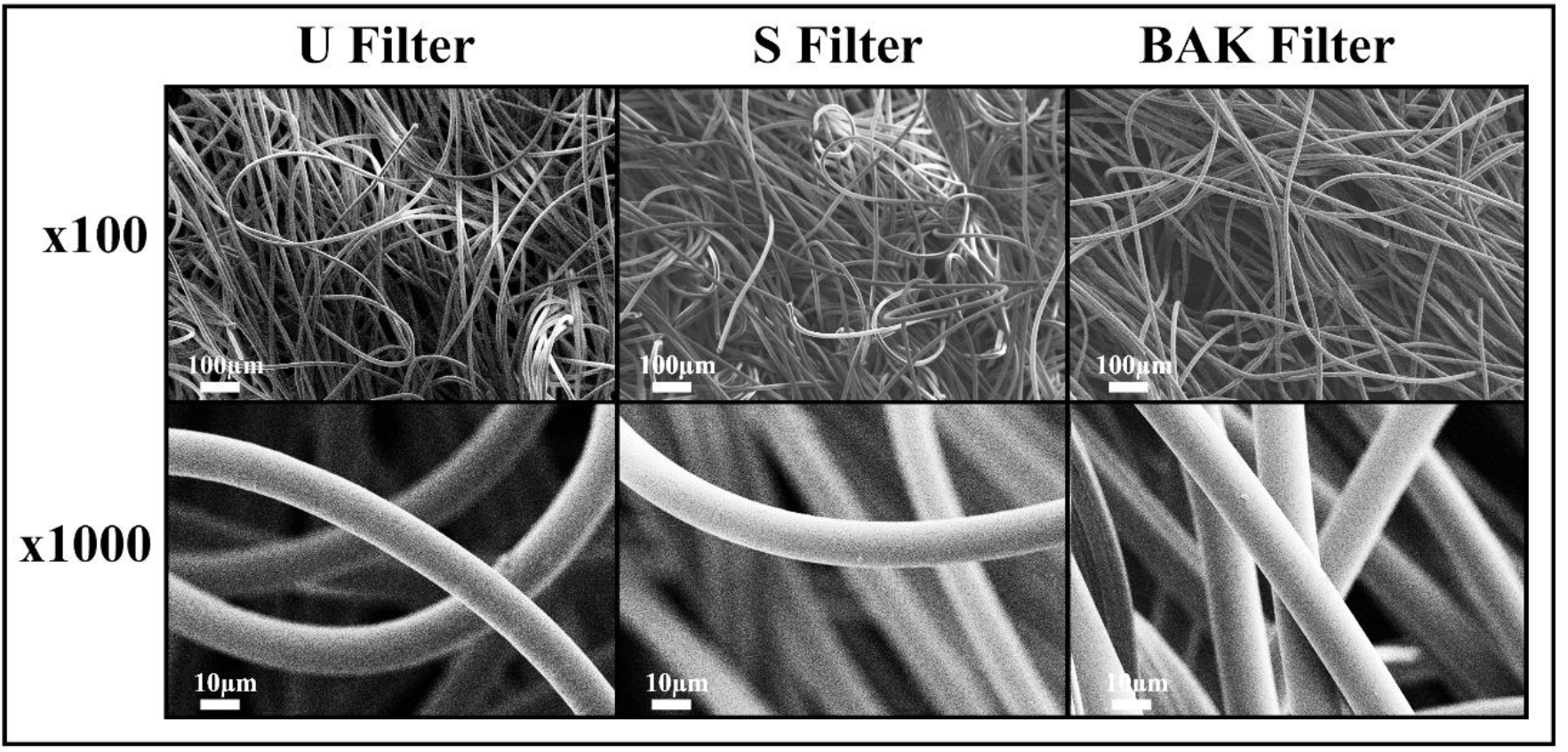
Morphology of the non-woven face mask filters by Field Emission Scanning Electron Microscopy. Untreated filter (U Filter), filter treated by dip coating with the ethanol-based solvent (S Filters) and filter with 0.46±0.13% *w/w* of biofunctional BAK coating (BAK Filter) at two magnifications (x100 and x1000).

FESEM observation showed no signs of porous morphological change after performing the dip coating with both ethanol-based solvent or the BAK compound. These results suggest no change of breathability nor bacterial filtration efficiency required for their commercialization according to the European standard Community Face Coverings CWA 17553:202.

### 3.3. Antiviral tests using the biosafe SARS-CoV-2 viral model

Phage phi 6 is a double-stranded RNA virus with three-part, segmented, totalling ~13.5 kb in length. Even though this type of lytic bacteriophage belongs to the Group III of the Baltimore classification^30^, it was proposed here as a viral model of SARS-CoV-2, due to biosafety reasons, as it has also a lipid membrane around their nucleocapsid. Thus, BAK Filters showed potent antiviral activity (100% of viral inhibition, see Figures 3 and 4). Bacterial lawns had clearly grown in the plate and no plaques were observed after 1, 10 or 30 minutes of contact between the BAK filter and the SARS-CoV-2 viral model. Furthermore, the U Filters and S Filters showed similar results to control of no antiviral activity (see Figures 3 and 4).

**Figure 3.**
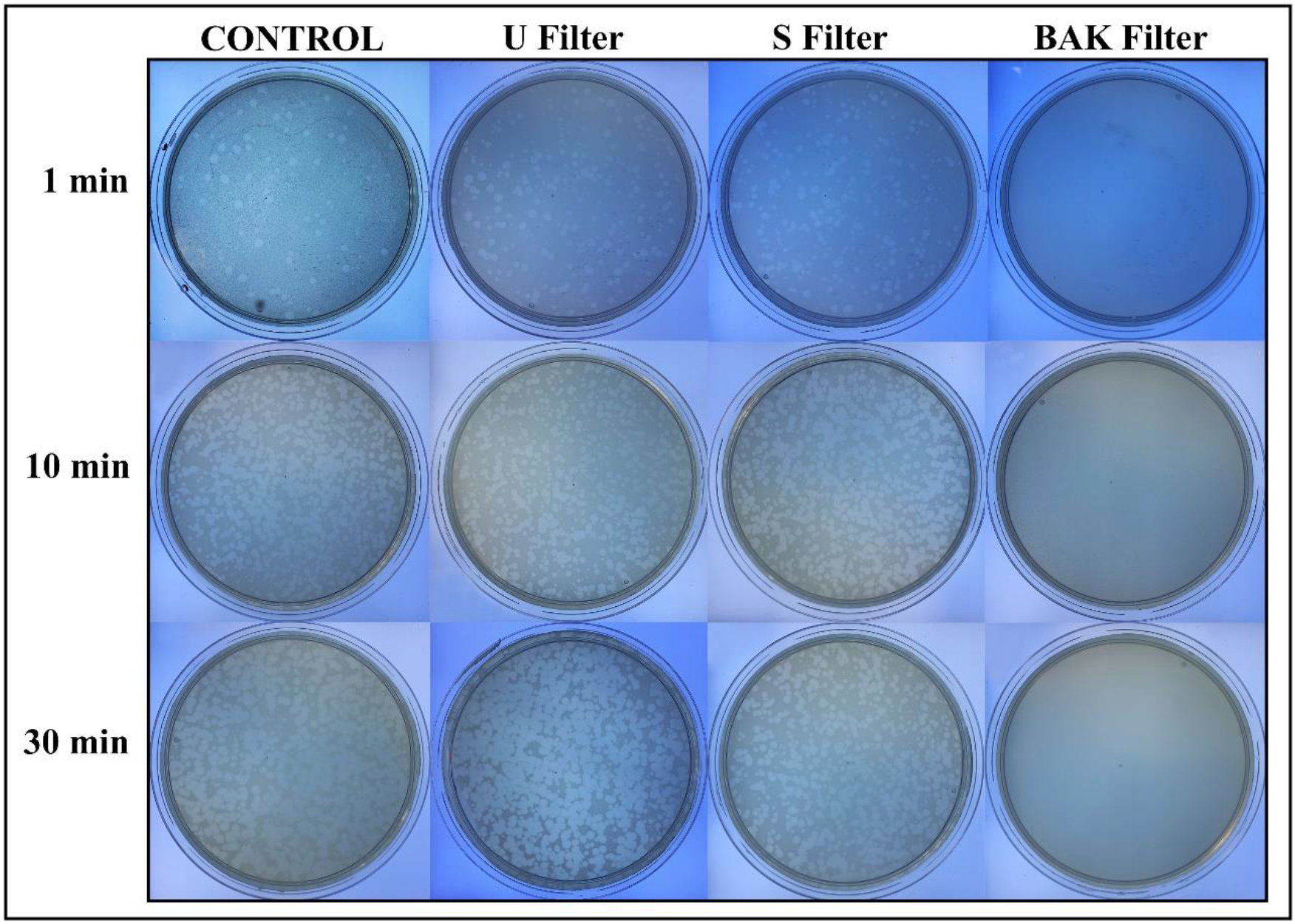
Loss of phage phi 6 viability measured by the double-layer method. Undiluted results for control, untreated filter (U Filter), filter treated by dip coating with the ethanol-based solvent (S Filters) and filter with the biofunctional BAK coating (BAK Filter) at 1, 10 and 30 minutes of viral contact.

The phage titers of each type of face mask filter sample were calculated and compared with the control (see Figure 4).

**Figure 4.**
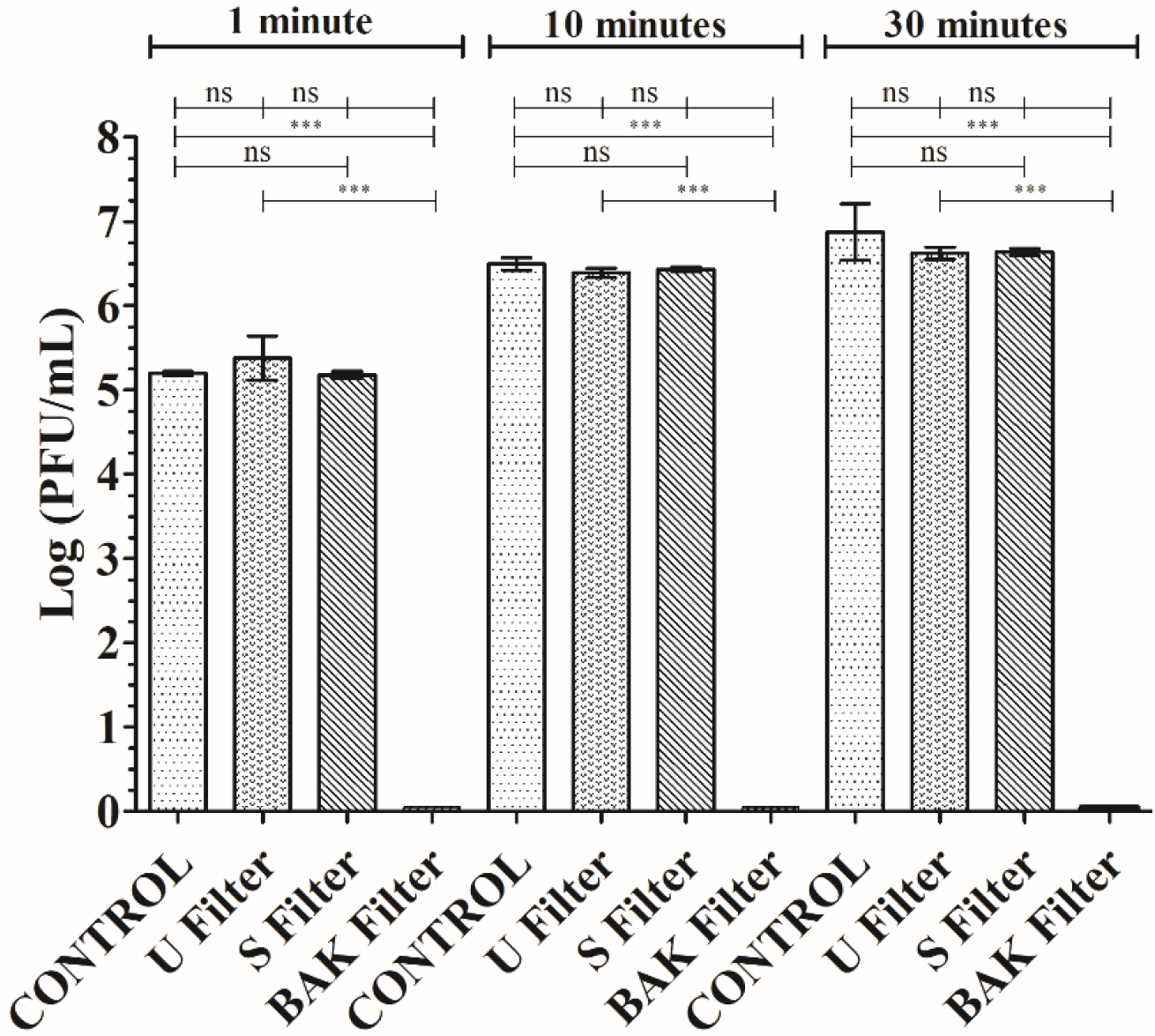
Titration after double-layer method with the phage phi6 viral model. Logarithm of plaque-forming units per mL (log(PFU/mL)) of the control, untreated filter (U Filter), filter treated by dip coating with the ethanol-based solvent (S Filters) and filter with the biofunctional BAK coating (BAK Filter) at 1, 10 and 30 minutes of viral contact.

Figure 4 shows that the titers obtained by contacting the phages with the U or S Filters are similar to the control. However, the BAK Filters displayed a strong phage inactivation.

### 3.4. Antiviral tests using SARS-CoV-2

The results achieved with the TCID50/mL method about the reduction of infectious titers of SARS-CoV-2 after 1 minute of contact with the control, the U Filters, the S Filters and the BAK Filters containing the biofunctional coating are shown in Figure 5.

**Figure 5.**
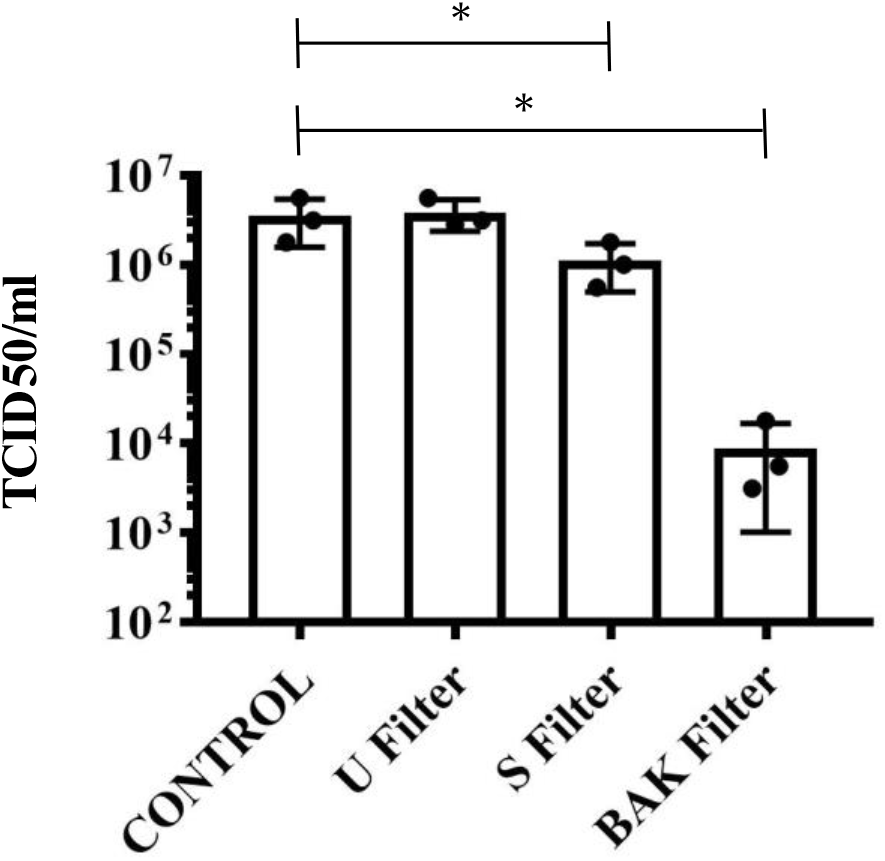
Reduction of infectious titers of SARS-CoV-2 after 1 minute of contact. Untreated filter (U Filter), filter treated with the ethanol solvent (S Filter), filter with the biofunctional BAK coating (BAK Filter) and control by the TCID50/mL method.

These results clearly demonstrate that the BAK Filter is very effective against SARS-CoV-2 even after 1 minute of contact. This is also in good agreement with the antiviral results of the biosafe viral model used in this study (see Figures 3 and 4).

### 3.5. Antibacterial tests

The antibacterial results of the treated filters against MRSA and MRSE multidrugresistant bacteria are shown in Figure 6.

**Figure 6.**
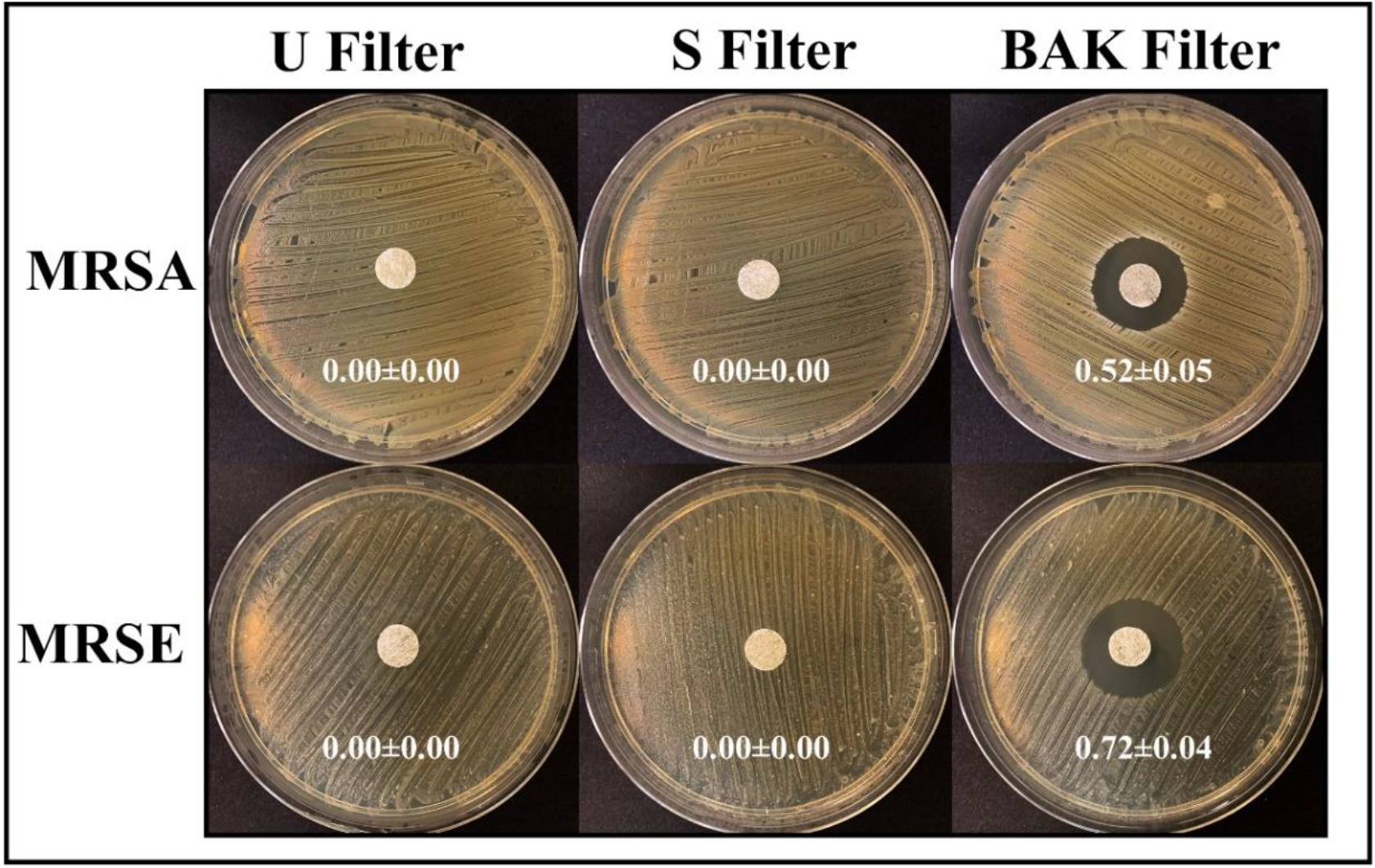
Antibacterial agar disk diffusion tests. Untreated filter (U Filter), filter treated by dip coating with the ethanol-based solvent (S Filters) and filter with the biofunctional BAK coating (BAK Filter) after 24 hours of culture at 37 °C. The normalized widths of the antibacterial “halos”, expressed as mean±standard deviation and calculated with equation (1), are shown in each image.

Filters treated by dip coating with 70% Ethyl alcohol containing 0.1% benzalkonium chloride showed high antibacterial activity against MRSA and MRSE, being even more effective against the last strain. The antimicrobial mode of action of quaternary ammonium compounds (QACs) such as BAK against both bacterial and viral phospholipid membranes are attributed to the positively charged nitrogen atoms. This cause eradication of surface bacteria and common viruses such as influenza by disrupting their phospholipid bilayer membrane^32^, the glycoproteinaceous envelope, and the associated spike glycoproteins interacting with the ACE2 receptor in the infection of host cells^52^. For this reason, BAK is extensively found in many household disinfecting wipes and sprays, and are also used as additives in various soaps and nonalcohol-based hand sanitizers^31,53,54^.

Here in this paper it is demonstrated a new face mask filter with antiviral and antibacterial properties against Gram-positive multidrug-resistant bacteria to reduce COVID-19 infections (from touching the filter masks and aerosol transmission in both senses). This face mask filter has been developed here by dip coating (a simple, low-cost, reliable and reproducible method) a commercial nonwoven filter where a thin coating of BAK were deposited onto the surface by physically adsorption^55^. The same biofunctional coating procedure could be applied to any type of face mask or biodegradable filters. This also represents a solution to the need for bio-based facemasks to counter coronavirus outbreaks^38^. The manufacturing procedure by dip coating with BAK opens up a broad range of applications that are demanding urgently new antimicrobial approaches. Thus, this technology may be also used for the fabrication of antimicrobial clothes, gloves, etc, for health personnel or to produce antimicrobial filters able to inactivate aerosols containing SARS-CoV-2 or Gram-positive multidrug-resistant bacteria in other applications. The antibacterial activity of these filters against MRSA and MRSE, and their viral inhibition capacity against SARS-CoV-2 and the enveloped phage phi 6 viral model demonstrates their broad-antipathogenic protection. Nonetheless, further research is required in order to ensure the safety use of the developed filters to combat the increasing COVID-19 spread.

## 4. Conclusions

Non-woven benzalkonium chloride-containing filters capable of inactivating SARS-CoV-2 and multidrug-resistant bacteria such as methicillin-resistant *Staphylococcus aureus* and *Staphylococcus epidermidis,* were developed here. This antimicrobial filter produced by a reproducible and economic procedure demonstrated excellent antiviral properties (100% viral inactivation already after 1 minute of contact) against the SARS-CoV-2 and the phage phi 6 phage viral model, which is an RNA enveloped virus like the SARS-CoV-2. Therefore, the developed antiviral filters can be used in face masks and thus are very promising to prevent the spread of SARS-CoV-2 in the present COVID-19 pandemic.

## Acknowledgments

The authors would like to express their gratitude to the Fundación Universidad Católica de Valencia San Vicente Mártir for their financial support through the Grant 2020-231- 001UCV (awarded to Á.S-A). This research was also supported by grants from the Japan Agency for Medical Research and Development (AMED) (20fk0108270h0001, 20fk0108263s0201). We would like to thank Dr. Yoshio Koyanagi and Dr. Kazuya Shimura (Kyoto University) for setup and operation of the BSL-3 laboratory.

